# Oxygen uptake in human donor corneas: the centripetal gradient and its changes in culture and correlation with wounds and aminophylline treatment

**DOI:** 10.1101/2025.06.26.661845

**Authors:** Ana M. Sandoval-Castellanos, Jeevan Mann, Brian Reid, Jennifer Y Li, Mark J Mannis, Min Zhao

## Abstract

Oxygen is a diatomic element abundant in the atmosphere and is essential for homeostatic and metabolic processes within the cornea. The avascular cornea uptakes oxygen from the atmosphere and performs refractory and protective functions that are essential for eyesight. However, the spatial-temporal profile of oxygen uptake (O_2_U) in human corneas remains uncharacterized in literature. Utilizing the micro optrode technique and donor corneas, we demonstrated that the limbus has a significantly higher O_2_U than the cornea center, exhibiting a pattern of centripetal gradient of O_2_U. This pattern is thus conserved in the corneas of mice, rats, non-human primates, and humans, representing a metabolic feature of physiological significance, which may have diagnostic and prognostic value. Being cultured at 37°C and 5% CO_2_, this centripetal pattern became more prominent in the first three days, and less so on day 4. The pattern remained in the corneas with injury, increased O_2_U appears to correlate with wound healing, similar to patterns observed in animal models with a dynamic increase of metabolism in the periphery, followed by the limbus, and subsequent return to baseline. Administration of the drug aminophylline prevented a decrease of O_2_U at day 4. We conclude that the centripetal pattern of O_2_U is conserved across species and in the human cornea. Combined, our results suggest the functional and clinical significance of the mesoscopic metabolism pattern for further study and the use of aminophylline as a therapeutic agent to increase O_2_U and enhance corneal wound healing.

## 1. Introduction

Oxygen is a diatomic element that in its molecular form (O_2_) is essential for aerobic life on Earth [1]. Molecular oxygen is readily abundant in the atmosphere at an approximate concentration of 21%. In the human physiological system, the lungs have adapted to facilitate the movement of oxygen into the bloodstream through alveolar tissue. The diffusion of oxygen from the bloodstream into tissues is dependent on several factors including pH, concentration of oxygen, temperature, and molecular effectors such as 2,3-bisphosphoglycerate [2, 3]

Oxygen is essential for the metabolic process of aerobic respiration, serving as the final acceptor of electrons from oxidative phosphorylation [4]. Furthermore, oxygen plays a key role in wound healing as a mediator of cellular processes such as collagen deposition, epithelialization, fibroplasia, and resistance to infection through its modulation of cellular signaling and transcription [5]. Reactive oxygen species (ROS), initially considered to be deleterious byproducts of metabolism, have been implicated in cellular signaling pathways and protein modification. Notably, recent studies have identified the importance of ROS in the facilitation of tissue regeneration through its participation in angiogenesis, cell migration, cell proliferation, and epithelialization [6, 7].

The cornea is one of the few virtually avascular tissues in the human body. Without abundant vascularization, the cornea relies on the diffusion of atmospheric oxygen through the tear film and nutrient absorption from the aqueous humor [8, 9]. Extended periods of hypoxia within the cornea can cause swelling, increased lactic acid production, reduced intracellular pH, and permanent morphological damage [10, 11]. Resultant morphological and physiological alterations of the cornea from these conditions lead to aberrant light scattering, affecting the quality of vision, highlighting the importance of cornea homeostatic maintenance [12].

Previous studies measured the oxygen uptake (O_2_U) at the human cornea, however, there was no mention of a specific region of the ocular surface [13, 14], or measurements on the limbus or conjunctiva [15], due to the probe size (>3mm in diameter). Using an advanced probe with size of 5 μm, our group has recently elucidated the O_2_U spatiotemporal profiles of *ex vivo* corneas from mice, rats, and rhesus monkeys, and mouse cornea *in vivo*, revealing a centripetal gradient of oxygen metabolism within *in vivo* and *ex vivo* [16]. In diabetes mellitus (DM) models previous studies have identified a dysregulation in the O_2_U, indicating a potential pathophysiological mechanism for corneas of animals affected by DM [13, 17, 18].

Significantly, the centripetal pattern of O_2_U 1) is lost in the diabetic cornea [17], 2) demonstrates a unique spatiotemporal and biphasic change in response to cornea injury [19] which is not present in wounded diabetic mice [18]. Although studies of human corneal O_2_U have been attempted, there has not been a spatio-temporal O_2_U characterization of the human cornea [14]. Therefore, the question we wanted to answer was if this centripetal gradient also exists in human corneas.

The administration of pharmacologic drugs, such as aminophylline, present the opportunity to enhance oxygen consumption, and possible wound healing in the cornea [20]. When aminophylline is applied to corneal wounds it enhances the endogenous electric current by increasing chloride and sodium fluxes, validating its potential role in enhancing wound healing and restoring native metabolic patterns to the eye [20, 21]. Thus, the next question was if aminophylline increased O_2_U at the human ocular surface.

Given the importance of oxygen in the process of cornea homeostasis and wound healing and the direct absorption of oxygen from the atmosphere, noninvasive methods of assessing oxygen consumption of the cornea remain highly clinically relevant. Numerous methods of quantifying oxygen flux are historically well established but have limitations such as the consumption of oxygen, interference, impassivity, and artifacts [19, 22]. Recent innovations in the field of oxygen sensors led to the development of the scanning micro-optrode technique (SMOT), allowing for an accurate, noninvasive, measurement of O_2_U. The SMOT operation is based on the principle of fluorescence quenching where an oxygen-sensitive fluorophore at the tip of the optrode is excited by blue-green light and is quenched by oxygen [16, 23]. The use of this unique technique allows for the safe and reliable spatiotemporal quantification of oxygen flux within tissues of interest. Furthermore, it poses a technical advantage due to the size of the probe, which is much smaller (∼ 5-10 μm) in comparison to a standard probe (∼ 3 mm), providing high spatial and temporal resolution.

Therefore, using the SMOT technique, we proposed to accurately and noninvasively characterize the O_2_U in human corneal tissues *ex vivo*. We measured O_2_U at the center, periphery, limbus, and conjunctiva of the ocular surface in donor human corneas for up to five days post*-*mortem (0 to day 4). We found a centripetal trend of O_2_U decreasing from the center of the cornea to the conjunctiva which was preserved throughout five days. On the last day, corneas were incubated with aminophylline, resulting in an increased O_2_U.

## 2. Methods

All procedures were approved by the Institutional Review Board (IRB) at the University of California, Davis (protocol #268125).

Oxygen flux measurements were performed through five days: from the day of arrival (denoted as day 0), then daily to the last day (day 4), a total of five days.

### 2.1 Human corneal tissue

Human donor corneas were obtained from Sierra Donor Services Eye Bank (Sacramento CA, US). Tissue was collected from 1-4 days post-mortem and shipped overnight immersed in a container with chilled optisol media (Figure 1A), inside a box with ice. The demographics of the donors were as follows: 75% female, 5% male; age 60 ± 4 years (Figure 1B). Upon arrival (and each following day until day 4), corneas were stained with fluorescein stain (I-Glo, JorVet, Cat no. J-1191) to obtain images of the condition/health of the epithelium.

**Figure 1.**
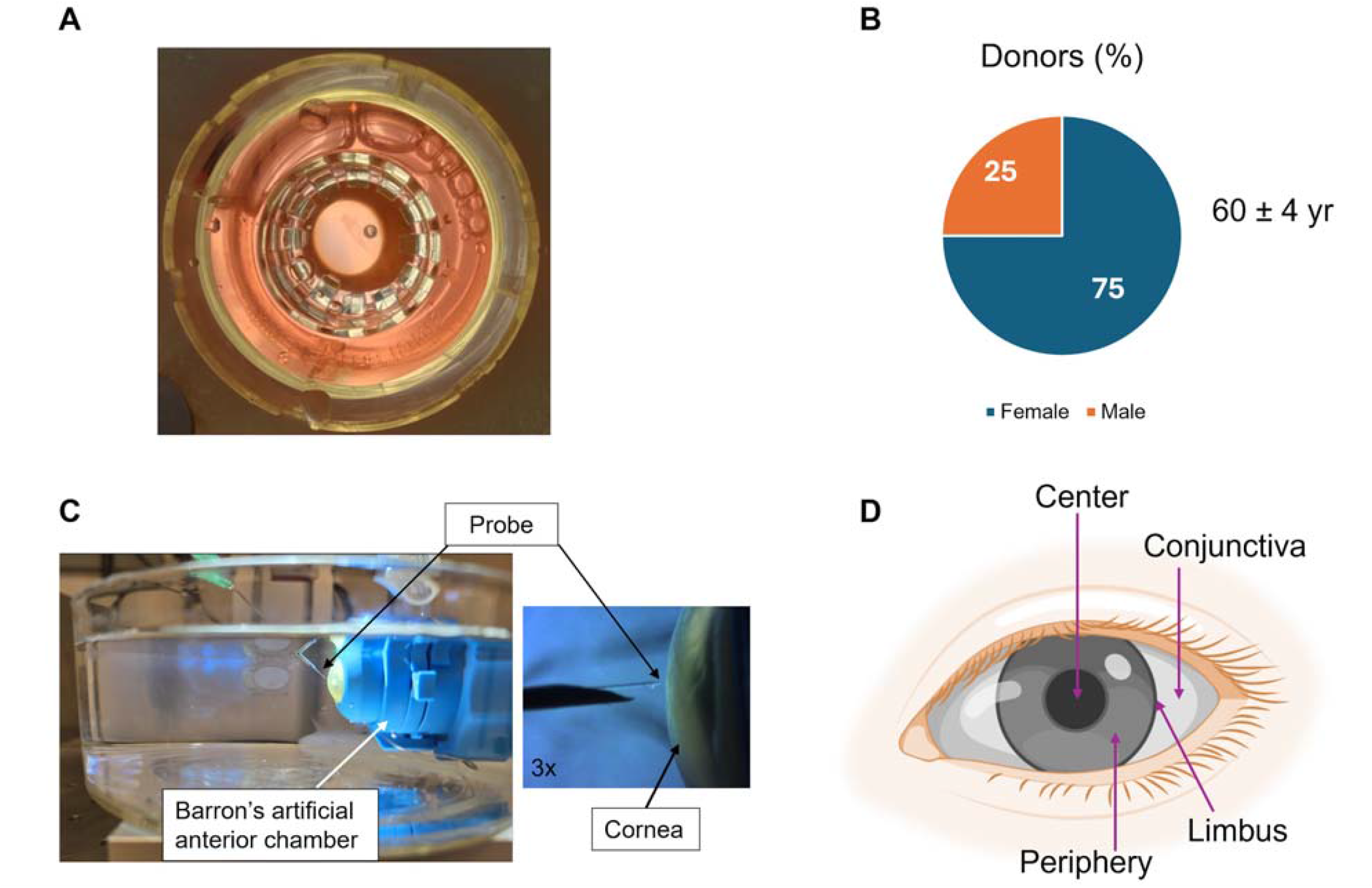
Experiment design. Donor human corneas were transported at 4°C in optisol medium (A). The demographics of the donors was: 75% female, 25% male, age 60±4 years old (B). Corneas were placed in a Barron artificial anterior chamber and immersed in artificial tear solution (BSS+). The oxygen probe was positioned at ∼ 10 μm from the cornea surface (C). Oxygen uptake (O_2_U) measurements were taken daily at the center, periphery, limbus and conjunctiva of the ocular surface (D).

Between measurements corneas were incubated at an air-liquid interface in supplemented serum-free culture medium at 37°C and 5% CO_2_ as described previously ([24]),) At the end of the study, at day 4, corneas were fixed with 10% normal formalin buffer (Globe Scientific, Inc., Cat no. 6518FL).

### 2.2 Wound size measurement

Wound size was measured with ImageJ (National Institutes of Health, version 1.54g). Using the fluorescein images and the freehand tool in the software, the total area of the cornea was determined. Then, the wound was measured (as it appeared green) and a wound size percentage was calculated relative to the cornea’s total area. This calculation was repeated throughout the 5 days of the experiment.

### 2.3 Scanning micro-optrode technique

The scanning micro-optrode technique (SMOT) and automated scanning electrode technique (ASET) software (version LV4), purchased from Applicable Electronics (New Haven, CT), have been previously described in detail [23]. Briefly, the SMOT is composed of a fiber-optic micro-needle probe (PreSens, Regensburg, Germany), an amplifier, a light source, and a software-controlled 3D micro-positioner. The probe is tipped with an oxygen-sensitive fluorophore. A fluorescent light source excites the fluorophore with blue-green light at 505 nm. Fluorescence quenching (rapid loss of fluorescence) occurs if oxygen is present. The rate of quenching is proportional to the oxygen concentration (pO_2_). Calibration of the probe in 0% and air-saturated (20.95%) dH_2_O is performed before and after measurements. The pO_2_ is calculated in real-time by the ASET software using the calibration values and oxygen flux calculated (using Fick’s first law of diffusion) with a Microsoft Excel spreadsheet template designed in-house.

### 2.4 Corneal oxygen uptake measurement

The cornea was mounted in a Barron artificial anterior chamber (Katena Products, Cat no. K20-2125) and immersed in artificial tear solution (BSS+ saline, Alcon Laboratory, Cat No.6508005. Figure 1C). The optrode sensor was carefully moved to ∼ 10 μm from the cornea surface. After waiting five minutes for the optrode signal to stabilize, the recording began, with the ASET software automatically moving the optrode 30 μm between the ‘near’ and ‘far’ positions at a frequency of 0.1 Hz, while simultaneously recording pO_2_ at the two positions. Then, O_2_ flux is calculated from these pO_2_ values using established equations [23]. Before and after measurements at the cornea surface, reference readings were taken more than 1 cm away from the specimen for 5 min to confirm the average zero O_2_ flux away from the cornea. O_2_ flux was measured at the following positions: center, periphery, limbus, and conjunctiva (Figure 1D). Timepoints were daily from day 0 (upon arrival) to day 4.

### 2.5 Aminophylline treatment

After the measurements on day 4, corneas were incubated at room temperature with 10 mM aminophylline (Santa Cruz Biotechnology, Cat no. sc-252368) dissolved in BSS+ for 20 min. Corneas were then measured again as above (section 2.4).

### 2.6 Data analysis

Data are shown as mean ± standard error of the mean (S.E.M.). Differences between samples were compared using Student’s t-test, and statistical significance accepted at 95% confidence limits (p < 0.05). All data analysis and statistics were done using Excel (Microsoft). Graphs were made using GraphPad Prism (version 10.4.1).

The methodology for calculating the lagged correlations between wound size and oxygen uptake (O_2_U) across days 0 to 4 involved a systematic statistical approach using Pearson’s correlation coefficient (r). For each specified lag period (e.g., day 0 to day 1, day 1 to day 2), data pairs were extracted from the human corneas (n=4), consisting of wound size percentage (%) values on the initial day and corresponding O_2_U values, at the center, periphery, and limbus, on the subsequent day. Pearson’s r was computed by first calculating the mean of each variable across the pairs, then determining the covariance as the sum of the products of the deviations of each pair from their respective means. The denominator was derived as the square root of the product of the sum of squared deviations for each variable, ensuring standardization. This process yielded correlation coefficients |r| ranging from 0 to 1, with values closer to 1 indicating that higher wound % predicted higher O_2_U.

## 3. Results

### Oxygen uptake in the human corneas has a centripetal gradient at day 0

After receiving the donor human corneas from the eye bank, we measured O_2_U at different positions: center, periphery, limbus, and conjunctiva. We found that O_2_U was different at each position, increasing from the center to the conjunctiva (Figure 2A), manifesting a centripetal pattern (Figure 2B). We observed significant differences between center and limbus, conjunctiva and center, and conjunctiva and periphery.

**Figure 2.**
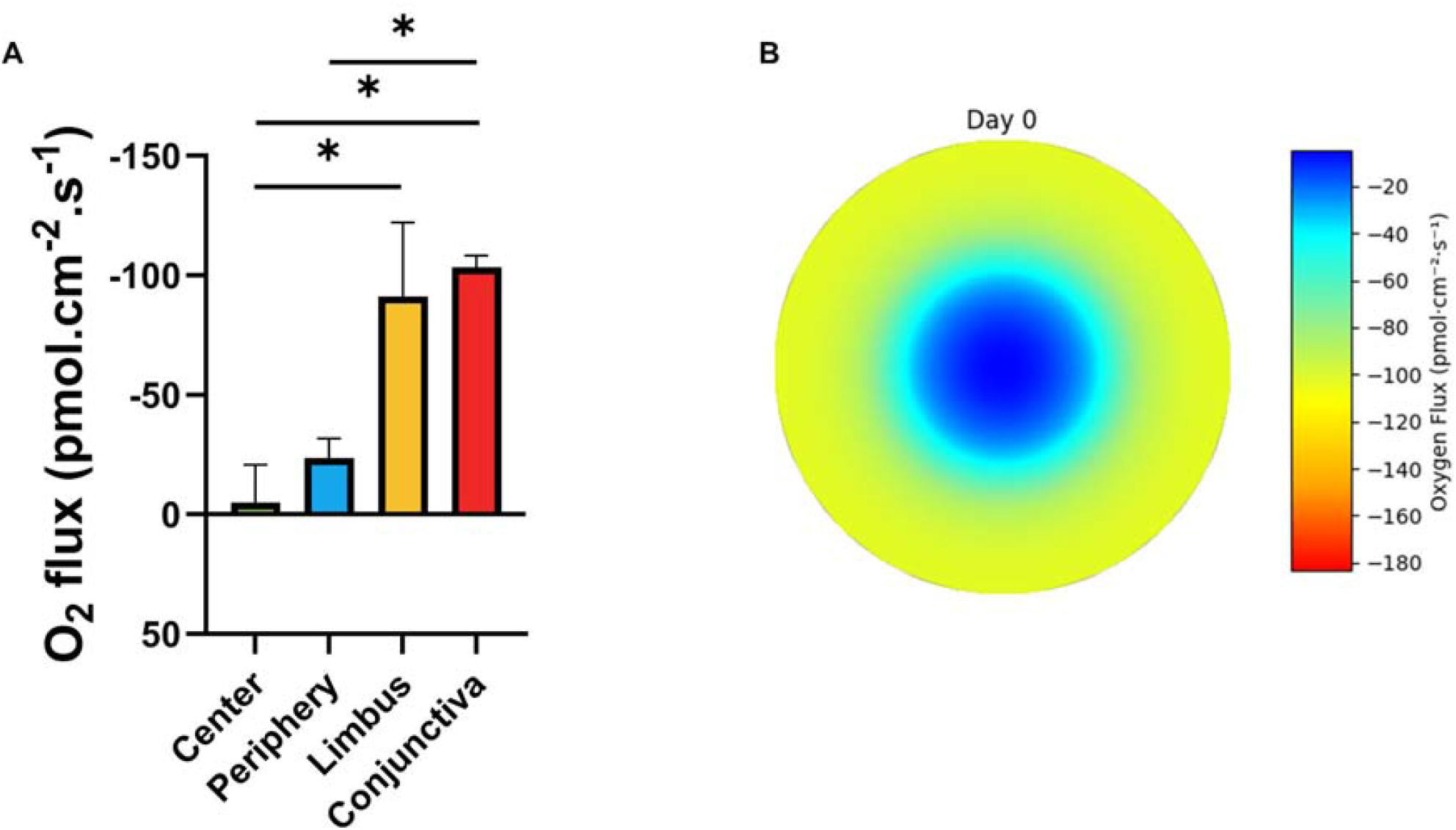
O_2_U has a centripetal trend at day 0. A) O_2_U is lower at the center of the cornea and higher at the limbus and conjunctiva. B) Representative heatmap of O_2_U in human donor corneas on day 0 showing a centripetal gradient across the ocular surface. Color scale bar shows color coded O_2_U values, where red indicates a high O_2_U, and blue a low O_2_U. Paired Student’s t-test, *p<0.05 was significant. n>4.

### O_2_U is dynamic at the ocular surface during 5 days of tissue culture

O_2_U was then measured up to day 4 at the center, periphery, and limbus (Figure 3). O_2_U showed dynamic changes at this position, increasing up to day 2, and then decreasing at days 3 and 4 (Figure 3A-C respectively). At the periphery, O_2_U on days 1 and 2 were significantly different from day 0 (p<0.02 and 0.03 respectively; Figure 3B). O_2_U at the conjunctiva was variable, increasing and decreasing without an obvious trend throughout this time period (Figure 3D). Heatmaps of O_2_U across the ocular surface show the dynamic changes from day 0 to 4 (Figure 3E).

**Figure 3.**
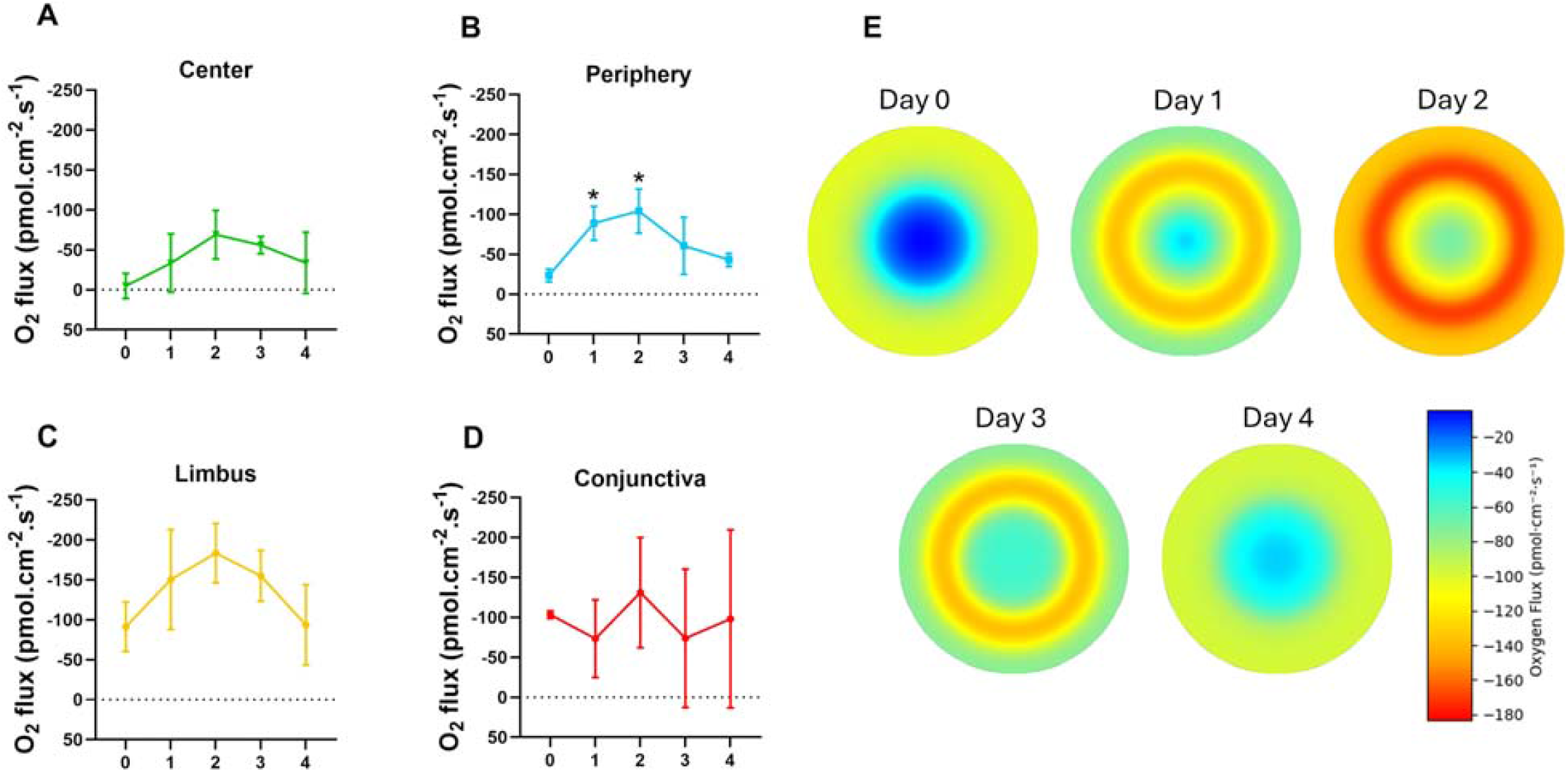
O_2_U at the ocular surface is dynamic during 5 days in culture. Human donor corneas were kept at 37 °C and 5% CO_2_, and oxygen flux was calculated at different positions: center (A), periphery (B), limbus (C), and conjunctiva (D). O_2_U increased, reaching a peak at day 2. Then, it decreased. For the conjunctiva, O_2_U was variable throughout this time (D). Heatmaps show dynamic changes of O_2_U across corneas over time (E). Blue = low O_2_U, red = high O_2_U. Unpaired Student’s t-test, *p<0.05 was significant. n>4.

### O_2_U changed during wound healing

Some corneas came with unintended wounds; we took advantage of that. We imaged the corneas with fluorescein dye from day 0 to day 4 to observe any changes in wound size. Figure 4A shows that the injury in the corneas either increased or decreased by day 4. We measured the injury size and calculated a percentage against the total area of the cornea. We found that wound size increased significantly in all corneas by day 1 (from 25 ± 10% to 56 ± 8%), which later decreased significantly by day 2 (33 ± 4%) and further went down at day 3 and maintained relatively the same size at day 4 (23 ± 20% and 23 ± 24% respectively; Figure 4B). From this data, it is seen that the peak of O_2_U on day 2 (Figure 4C) correlated with the time when significant wound reduction was observed. Additionally, O_2_U at the limbus was significantly higher in comparison to the center on days 1, 2, and 3 (Figure 4C).

**Figure 4.**
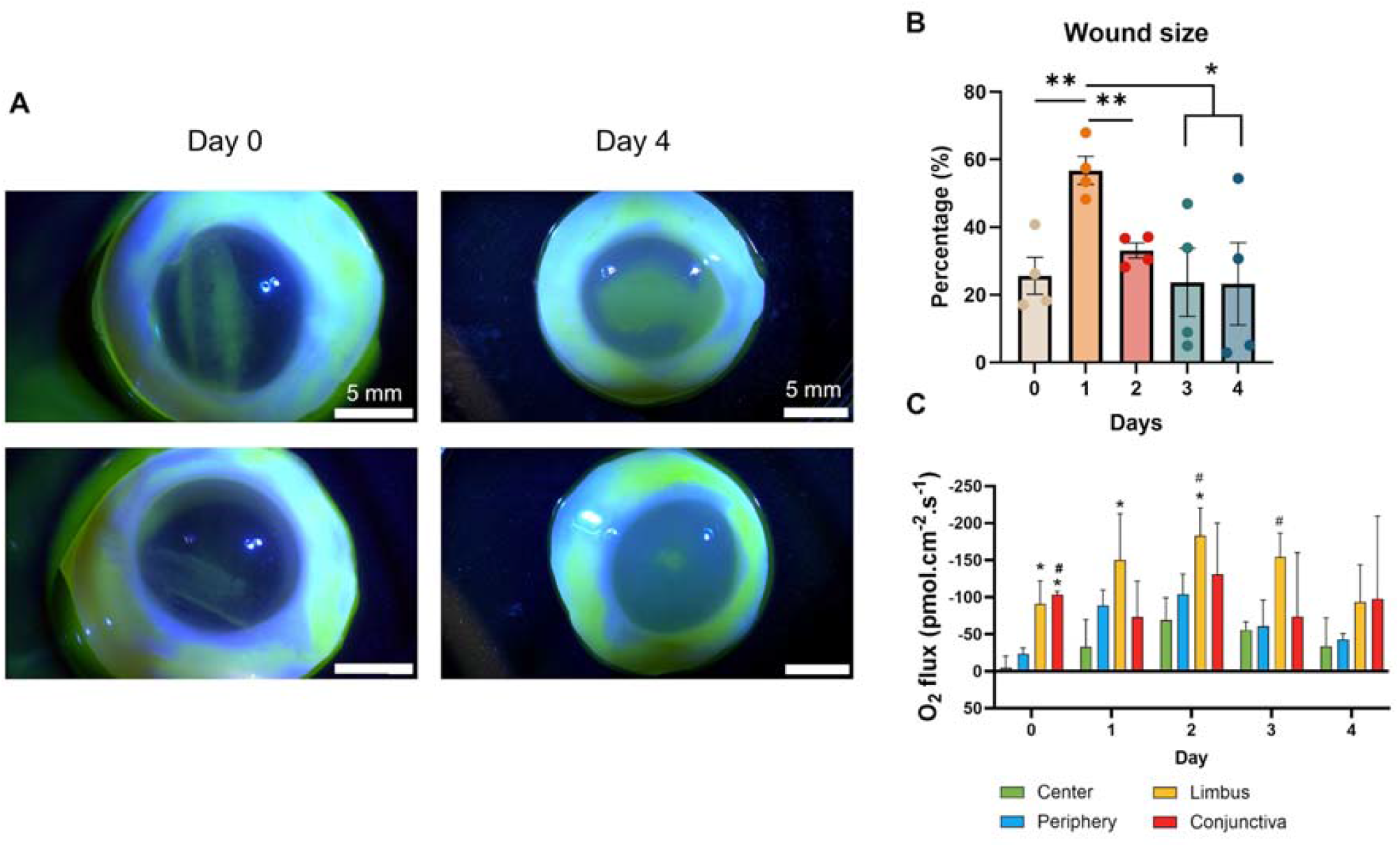
O_2_U changed during wound healing in human donor corneas. Corneas that had an injury upon arrival were used for the wound healing observation (green, fluorescent stain in A) and O_2_U measurement, and the injury size either decreased or increased after 4 days in culture (A, B). O_2_U was higher on day 2, when wound size drastically decreased (B, C). Then, as the wound size kept decreasing, oxygen flux was also lower. Paired Student’s t-test, n=4. Compared to the center: *p < 0.05, ** p<0.01. Compared to the periphery #p<0.05.

### Aminophylline increased O_2_U in human donor corneas

After oxygen measurement on day 4, we exposed the human cornea to 10 mM aminophylline for 20 min and measured oxygen at the center, periphery, limbus, and conjunctiva (Figure 5). We found that O_2_U increased after aminophylline incubation, from −33 ± 93 to −79 ±69 in the center, from −43±16 to −127 ± 64 at the periphery, from −93 ± 123 to −152±120 at the limbus and from −98 ± 222 to −166 ± 73 at the conjunctiva (units in pmol.cm^-2^.s^-1^). We observed that the centripetal trend on day 4 was conserved after aminophylline exposure. However, after aminophylline treatment, only O_2_U at the periphery was significantly higher compared to the center (Figure 5B).

**Figure 5.**
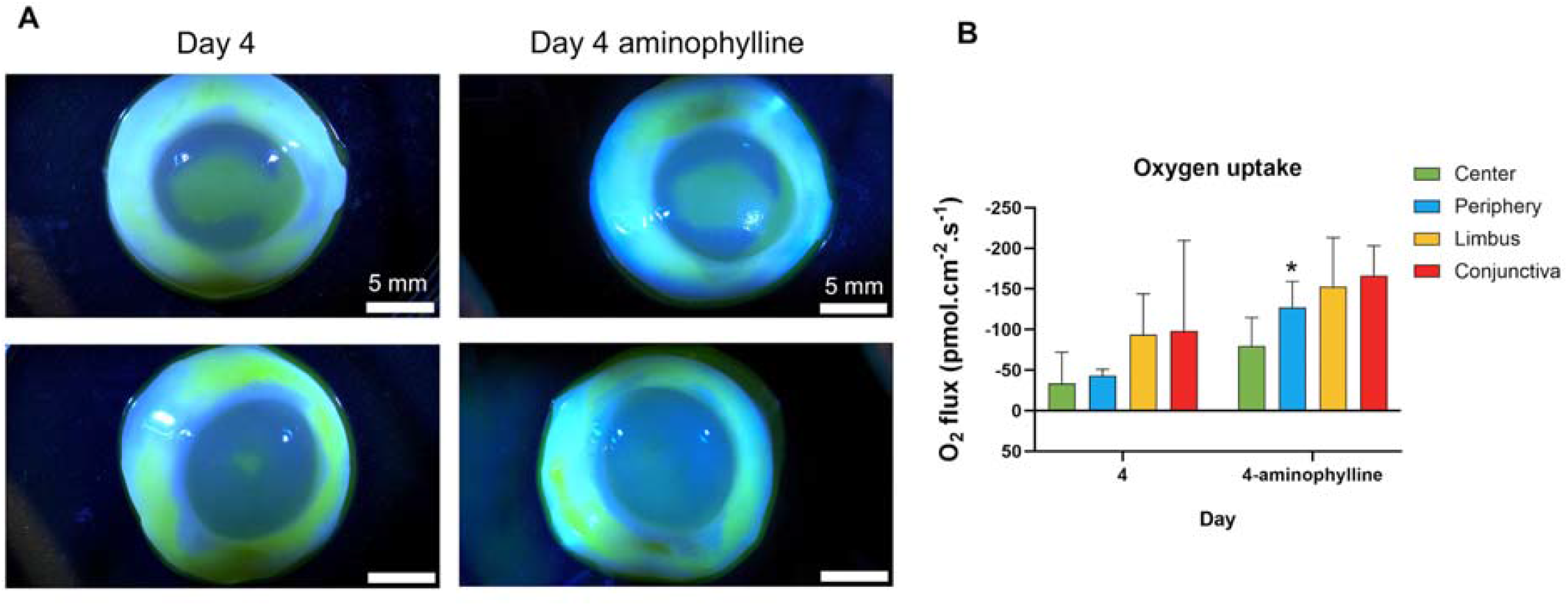
Aminophylline stimulated increased O_2_U in human donor corneas. A) Wounds can be observed before and after aminophylline treatment, suggesting that the injury in one cornea was reduced while it increased in the other cornea. B) O_2_U increased in all regions of the human cornea after aminophylline treatment. Paired Student’s t-test, n=4. Compared to the center: *p < 0.05.

## 4. Discussion

**Human donor cornea manifests a unique centripetal O**_**2**_**U**, lowest at the center and highest at the limbus and conjunctiva (Figures 2, 3). This unique and previously unreported centripetal pattern of O_2_U in human corneas is highly similar to the spatiotemporal pattern observed at the intact ocular surface in mice and rats, both *ex vivo* and *in vivo*, and non-human primates *ex vivo* [16, 17, 19]. Such a gradient pattern implies a difference in tissue-level metabolism. From existing literature and our observations of epithelial health during culture, the maintenance of mammalian corneal metabolism is reliant on the centripetal gradient, indicating conservation of centripetal corneal metabolism across different species of mammals [16, 19].

The human cornea is avascular (except the small limbal vascular arcade, terminal blood vessels located in the outer limbus [25-27]), therefore, O_2_U from the atmosphere is essential for its maintenance of metabolic homeostasis and for the proper functioning of corneal tissue [4, 11]. Oxygen flux is highest in the limbal region of the cornea, where the limbal epithelial stem cells (LESC) reside [28]. As a reservoir of LESCs, the limbus is functionally separated into two subpopulations: the inner limbus and the outer limbus. The outer limbus is primarily composed of quiescent LESCs, while the outer limbus contains active stem cells [29]. The active LESC population likely has an increased metabolic demand for oxygen due to their increased cell division and migration across the cornea to maintain homeostasis and heal wounds [16, 29-31].

Understanding the spatial temporal metabolic dynamics of the inner and the outer limbus could provide invaluable insights into the activation and proliferation of the LESC population within the two areas of the limbus. Particularly, it would be relevant to discern the metabolic differences in the limbal regions. Further spatial resolution at this level could resolve whether the cause of the deficit is from activation or proliferation defects.

Changes in the centripetal O_2_U pattern may indicate a significant metabolic aberration in diseases because this pattern is significantly compromised in diabetic corneas [13]. Aberration in this pattern thus may indicate underlying tissue-level metabolisms and are likely of clinical value, e.g., diagnostic and prognostic.

**O_2_U in the human cornea was dynamic through culture.** There was a trend in O_2_U throughout the five days in culture, a steady increase of O_2_U on days 1, 2, and 3, followed by a decrease on day 4, but still higher than that on day 0 (Figure 3). This is likely due to increased metabolism when the corneas were cultured in a medium with various nutrients, and kept at 37°C, suggesting viability and resilience of the tissues.

**O_2_U in the human cornea was dynamic through wound healing.** Wounding the cornea was not made on purpose. We speculate that part of the cornea surface was damaged by prolonged exposure to the air sometime between the passing of the patient and harvesting the corneas. This presented a unique opportunity to measure O_2_U trends during wound healing. Although not controlled or precise wounds, our data provided preliminary insights into O_2_U during wound healing. Upon arrival, and daily until day 4, fluorescein stain was used in the corneas to image the wounded area (Figure 4A). We found that wound size increased from day 0 to day 1, which then decreased (Figure 4B). This pattern was also observed in the O_2_U measurements. It is interesting to note that the increased O_2_U on day 2 happened at the same time as wound size decreased, highlighting that higher O_2_U is key for wound reduction in human corneas.

Cornea wound healing and regeneration is divided into two distinct stages: the initial latent phase induces intercellular and intracellular reorganization to facilitate the migration of epithelial cells; while the second stage, closure, can be described as cell migration, proliferation, differentiation, and stratification of cells [30]. The healing capacity of the cornea depends on the LESCs, located at the limbus. During wound healing, LESCs increase their rate of proliferation by up to 9-fold in the limbal region and about 2-fold in the central and peripheral regions [32]. A primitive population (quiescent) and active population of LESCs have been proposed [28, 29]. Transformation of the quiescent stem cell population into an active state increases their metabolic oxygen demand, and subsequent O2 consumption by the peripheral and central epithelium suggested increased healing responses required more oxygen. Rat corneal wounds *in vivo* showed a similar change in O_2_U of increased metabolism in the peripheral region and limbus following wounding [17]. These results suggest that, following injury, LESCs increase proliferation and migration thus higher metabolic demand for oxygen [33, 34].

Therefore, we proceeded with a Pearson correlation analysis to determine any correlation between O_2_U and wound size. The lag analysis of O_2_U and wound size percentage across days 0 to 4 provided a comprehensive view of healing dynamics in human corneas (Figure 6.). From day 0 to day 1, strong correlations (|r| = 0.70 for the center, 0.83 for the periphery, 0.94 for the limbus) showed that higher initial wound size % (e.g., 17.09 to 40.84 on day 0) predicts increased O_2_U (e.g., −146.159 to −396.305 on day 1), indicating an immediate metabolic response to wounding. From day 1 to day 2, with wound size peaking (e.g., 57.50 to 67.94), correlations weaken (|r| = 0.32 for periphery, 0.37 for limbus), reflecting varied O_2_U responses (e.g., −134.05 to −243.6) as wounds stabilize. From day 2 to day 3, as wound size % decreases (e.g., 36.68 to 46.99), correlations strengthen (|r| = 0.84 for center, 0.18 for periphery, 0.94 for limbus), with higher O_2_U (e.g., −95.102 to −287.259) suggesting continued healing. Finally, from day 3 to day 4, with wound size further decreasing (e.g., 8.96 to 46.98), moderate to strong correlations (|r| = 0.38 for center, 0.36 for periphery, 0.54 for limbus) indicate a sustained change in the oxygen demand, particularly at the limbus, highlighting a consistent relationship throughout the healing process. This analysis suggests a physiological context where a wound heals and changes in O_2_U during the healing process.

**Figure 6.**
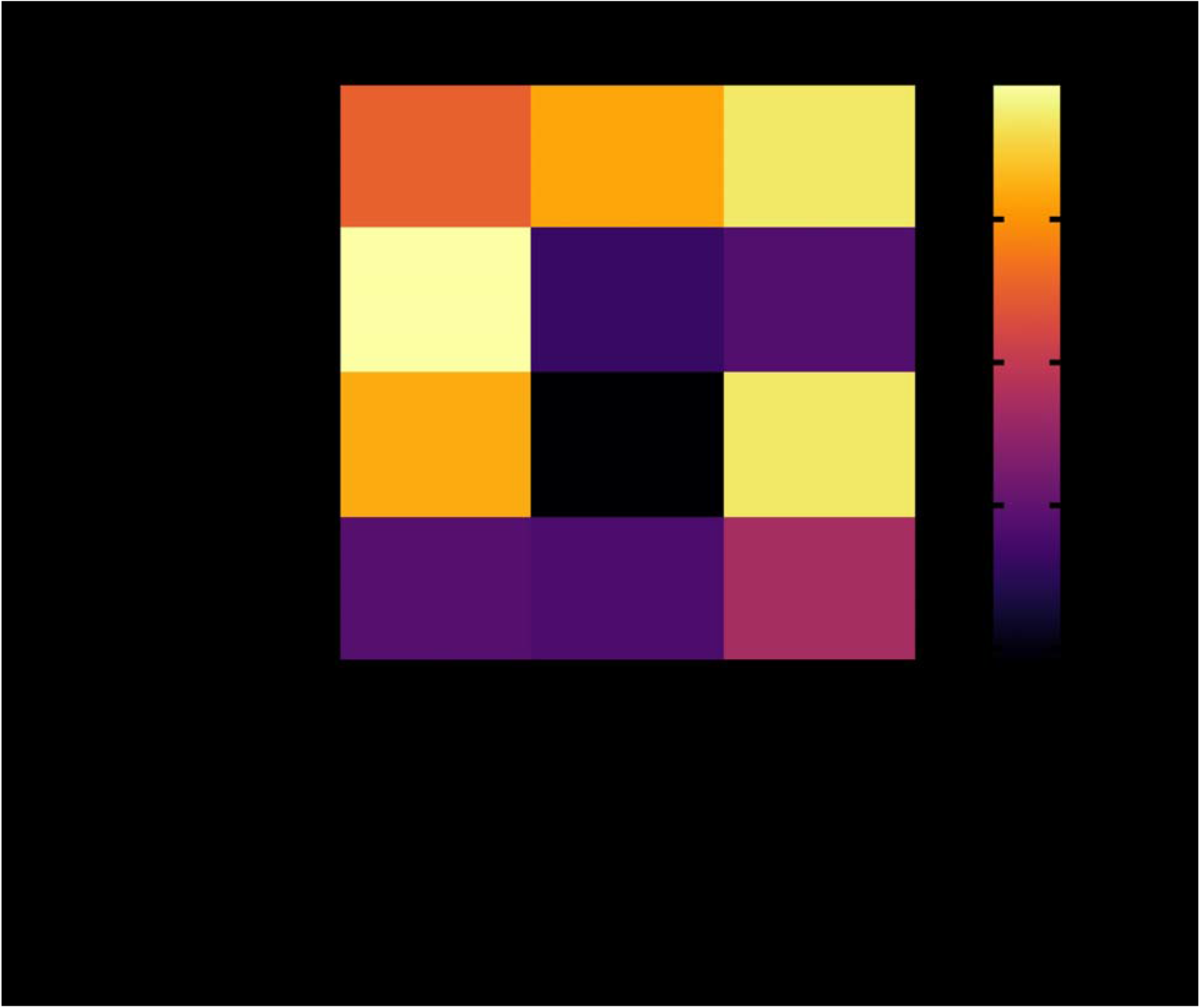
Correlation of wound healing and O_2_U during the healing process. This heatmap illustrates the relationship between wound size percentage (Wound %) and oxygen uptake (O_2_U) across four lag periods (day 0 to 1, day 1 to 2, day 2 to 3, and day 3 to 4). Pearson correlation analysis was employed to quantify this relationship, where each correlation coefficient (r) was calculated by comparing Wound % on the initial day of the lag (e.g., day 0) with O_2_U (center, periphery, limbus) on the subsequent day (e.g., day 1). The original r values range from −1 to 1, with negative values indicating that higher Wound % predicts higher O_2_U (reflected as more negative O_2_U values, indicating increased O_2_U due to metabolic demand during healing). The heatmap displays the absolute value of these correlations (|r|), scaled from 0 (weak or no relationship, dark purple) to 1 (very strong relationship, bright yellow), to emphasize the strength of the association regardless of direction. This pattern highlights that wound severity drives O_2_U dynamically, with peak effects in the early and mid-healing phases. n=4.

Aminophylline, a drug affecting ion channel activity, has been suggested to enhance the healing of corneal wounds, presumably by increasing electric current, specifically, stimulating increased chloride influx and sodium efflux [20, 21, 35]. After oxygen flux measurements on day 4, we exposed the human corneas to aminophylline for 20 minutes, then measured O_2_U. Aminophylline increased O_2_U, in all regions of the ocular surface (Fig. 5), suggesting aminophylline’s effects on aerobic metabolism of the cornea. Further experiments are required to understand whether the effect of aminophylline is additive and its ideal concentration.

## 5. Conclusion

Spatiotemporal characterization of the human cornea reveals a distinct pattern of centripetal O_2_U, greater O_2_U in the limbal region, compared to the center of the cornea. Spatial-temporal O_2_U characterization over 5 days showed distinct patterns suggesting an increasing metabolism. This gradient of O_2_U originating from the limbus restores oxygen flux within the center region, suggesting an underlying mechanism for corneal maintenance. Animal and human corneal tissues have conserved O_2_U metabolic patterns in wound healing. Aminophylline increased O_2_U in the cornea. SMOT offers a powerful tool to investigate tissue-level metabolism of corneas in health and diseases, including aging, diabetes mellitus, dry eye, and other diseases and injuries. Such mesoscopic spatiotemporal metabolic patterns and their responses to injuries and diseases provide a practical assessment of important pathophysiological conditions of the cornea.

## Acknowledgments

Sierra Donor Services Eye Bank for their timely response and shipping of human donor corneas when requested.

Suyash N. Goel for his help and support in making the heat map in Figure 2B and 3D. All members of the Zhao lab.

The Zhao lab thanks the Burns family, Mr and Mrs Meyers, and H. Schroeter for their generous donations that bolster research efforts.

